# Ultra-structural analysis of mineralized extracellular matrix in osteogenic monolayers and spheroids: comparison of sample preparation methods

**DOI:** 10.64898/2026.07.03.736266

**Authors:** Diamante Boscaro, Ursula Ludacka, Pawel Sikorski

## Abstract

Accurate evaluation of extracellular matrix (ECM) mineralization at the nano-scale is essential for establishing relevant *in vitro* bone models. This is particularly important with the development and increased application of three-dimensional (3D) cell models for biological research. Transmission electron microscopy (TEM) allows to perform ultra-structural analysis of cells and ECM organization, but its application in *in vitro* bone models remains limited, due to the potential alteration or loss of the mineral phase during sample preparation. In this study, we compared two TEM sample preparation methods the conventional chemical fixation and the anhydrous methods to evaluate their ability to preserve the mineralized ECM in MC3T3-E1 cells cultured as monolayers and as alginate-encapsulated bone spheroids. Chemical fixation preserved cellular ultra-structure and collagen organization, allowing for detailed assessment of cells and ECM organization. Although mineral deposits were detected and their needle-like morphology assessed, characterization of more immature deposits was partially limited by the effects of uranyl acetate and the overall sample preparation process, which could lead to alteration or loss of less stable mineral phases. The anhydrous preparation method resulted in limited preservation of cellular and ECM morphology and did not allow reliable identification of mineral deposits. When applied to spheroids, the chemical fixation method preserved the 3D architecture, collagen-rich ECM and inner mineral deposits, confirming spheroids as a relevant model for bone studies. Overall, these results highlight the need for optimized sample preparation strategies that preserve both ultra-structure and mineral components for accurate nano-scale characterization of bone mineralization.

## INTRODUCTION

Understanding the organization and mineralization of the extracellular matrix (ECM) on the nano-scale is fundamental in bone tissue engineering studies, especially when investigating the formation and deposition of the mineralized matrix. Bone ECM is a highly organized environment, composed of an organic part (40%), made mostly of type I collagen, and an inorganic part (60%), consisting of calcium-deficient apatite (1, 2). One of the most used cell lines to study bone mineralization is the murine pre-osteoblastic MC3T3-E1 cell line, which can be differentiated into mature osteoblasts in the presence of ascorbic acid and β-glycerophosphate, leading to the formation of a highly collagenous and mineralized ECM (3–5).

Although this cell line has been extensively characterized with regard to matrix deposition, most of these studies have been conducted on traditional two-dimensional (2D) cell cultures. However, it is now well established that 2D cell models cannot fully recapitulate physiological and pathological conditions observed *in vivo*. To overcome these limitations and develop more physiologically relevant *in vitro* systems, three-dimensional (3D) cell culture models have been introduced. The rapid advancement of 3D culture models, such as spheroids, organoids and organ-on-a-chip, (6), has created a growing need not only to evaluate cellular behavior within these complex microenvironments but also to reassess and adapt established characterization techniques for their use in 3D models (7, 8).

In bone studies, commonly used techniques to assess mineral deposition are based on colorimetric or fluorescent dyes (9). However, these techniques present several limitations. First, they lack specificity for bone mineral, as most dyes bind to calcium ions, which can lead to false-positive results. Their application in 3D culture systems is also limited, as the literature reports inconsistent results regarding staining efficiency and penetration in these models (10–13). Furthermore, these methods do not allow for a comprehensive evaluation of ECM ultrastructure, as collagen fibril organization and the general matrix architecture cannot be visualized thoroughly.

To overcome these limitations, high-resolution imaging techniques that allow simultaneous evaluation of mineral deposition and ECM ultrastructure are required. In this context, transmission electron microscopy (TEM) is a powerful tool, as it allows direct visualization of collagen fibril organization, overall matrix architecture, and mineral deposits at high resolution. Importantly, for 3D cell models such as spheroids, TEM allows comprehensive assessment of the entire construct, including cellular morphology, mineral localization, and ECM organization (14). Moreover, in systems where spheroids are embedded within a supporting biomaterial, TEM enables evaluation of potential structural interac-tions between the cells, the formed matrix and the surrounding hydrogel. The possibility of performing elemental analysis, such as Energy Dispersive X-ray Spectroscopy (EDX), provides an additional advantage for mineral deposition studies, as it enables precise identification of the elements present in the sample. Despite its high spatial resolution and ability to reveal ultra-structural details, the application of TEM for the analysis of 3D bone cell models remains limited (15) and is often not included in mineralization assessment workflows. TEM is in fact most often applied to native bone tissue sections and 2D cell cultures (16–18). In addition, the application of TEM to mineralized *in vitro* models presents some methodological challenges, mostly related to sample preparation. The traditional chemical fixation method, while being effective in preserving cellular and ECM ultrastructure, may alter or partially dissolve mineral phases, affecting their morphology or distribution. Furthermore, the use of uranyl acetate both during sample preparation and for contrasting may also influence mineral preservation, as, due to its acidic nature, can interact with phosphate groups and potentially contribute to the partial dissolution or modification of immature calcium phosphate deposits. Although anhydrous processing methods have been proposed to improve mineral preservation, they remain relatively underutilized in studies assessing mineral deposition in *in vitro* bone models, as the ultrastructure of the cells is not well preserved (19–22).

In this study, TEM was applied to both monolayer cell cultures and alginate-encapsulated spheroids to perform an ultra-structural characterization of the cells and their deposited ECM. Two different sample preparation methods were used on monolayer cell cultures, to evaluate which approach provides optimal preservation of the mineral structures. The method that better preserved the deposited ECM was used for spheroids analysis. In addition to comparing the different sample preparation methods, both contrasted and non-contrasted samples prepared with the same method were imaged to assess the effect of contrasting aqueous solutions on the deposited mineral.

## MATERIAL AND METHODS

### Cell culture

MC3T3-E1 subclone 4 (ATCC, CRL-2593) were cultured in standard conditions at 37 °C and 5% CO_2_ in minimum essential alpha medium (MEM-α without ascorbic acid; ThermoFisher, A1049001) supplemented with 10% fetal bovine serum (FBS; Sigma-Aldrich). To induce osteogenic differentiation, the growth medium was supplemented with 50 µg*/*mL of L-ascorbic acid 2-phosphate sesquimagnesium salt hydrate (Sigma-Aldrich, A8960) and 5 mM of β-glycerophosphate disodium salt pentahydrate (Sigma-Aldrich).

### Spheroids formation

Spheroids were obtained using the micro-mold technique (23). Briefly, 1.5% agarose (SigmaAldrich) molds were made from #24-96 silicon molds (MicroTissues 3D Petri Dish micro-mold spheroids) and were placed in a 24-well plate. The cells were collected and re-suspended in 1 mL of RM, and 75 µL of cell suspension were added to each mold, followed by 1 mL of RM added into the well. After overnight incubation, the molds were placed upside down in a new 24-well plate and centrifuged to allow spheroids collection. The molds were removed and the media with spheroids was collected in a 15 mL tube and centrifuged. The sedimented spheroids were retrieved and resuspended in a solution made of an equal volume of RM and sterile filtered 2% Alginate (Alg) solution (G fraction: 0.68, GG fraction: 0.57, Molecular weight: 250000 g*/*mol, [Novamatrix (Sandvika, Norway)]). 150 µL of the spheroids solution was added into glassed-bottomed Petri Dishes (35 mm Dish with 14 mm bottom well, #1.5 glass – 0.16 - 0.19, Cellvis, Cat. #D35-10-1.5-N), covered with a GN-6 Metricel® 0.45 µm-47 mm sterile membrane and gelation was induced by adding 50 mM CaCl_2_ on top of the membrane for 5 minutes. After gelation, both the CaCl_2_ solution and the membrane were removed. The alginate disks containing spheroids were kept in 2 mL of media. Media was changed every second or third day. Spheroids were maintained with the same standard conditions as the monolayer cell cultures. Osteogenic media (OM) was used to induce osteogenic differentiation of the spheroids in the alginate disks.

### Transmission Electron Microscopy (TEM) sample preparation

Two TEM sample preparation protocols were used depending on the structural features of interest. Traditional chemical fixation protocol was employed on monolayer cell cultures and on spheroids for analysis of the cells and/or spheroids structure and overall ECM organization. The anhydrous protocol was employed on monolayer cell cultures to improve the preservation and visualization of the mineral deposits. Monolayer cells, for both sample preparation methods, were cultured on an Aclar film in a 12 well plate, while spheroids were maintained in standard conditions. All samples were cultured in both RM and OM. For the traditional chemical fixation method, monolayer cells were cultured for 2 and 3 weeks in RM and OM, while spheroids were cultured for 5 and 6 weeks. Both types of samples were fixed in 2% formaldehyde, 2.5% glutaraldehyde, and 0.025% CaCl_2_ in 0.1 M cacodylate buffer (pH = 7.4) overnight at room temperature (RT). The samples were washed with 0.1 M cacodylate buffer with 0.025% CaCl_2_ twice for 15 minutes. At this stage, the alginate disk containing the spheroids was cut into smaller pieces to facilitate chemical penetration during subsequent sample preparation. Samples were then post-fixed in 2% osmium tetroxide, 0.025% CaCl_2_ and 1.5% potassium ferrocyanide for 45 minutes. The samples were washed again with the same washing solution as mentioned before and dehydrated in increasing ethanol series (50%, 70%, 90%) for 5 minutes each. The samples were followed by uranyl staining (4% Uranyl Acetate (UA) in 50% ethanol) for 1 hour and 30 minutes. Lastly, the samples were washed first in absolute ethanol and then in acetone and embedded in a pure acetone and epoxy (AGAR) resin mix in a ratio of 2:1, 1:1 and 1:2 for 45 minutes each. The samples were then incubated in epoxy resin overnight on a rotator. The samples were embedded in fresh resin and polymerized at 60 °C overnight. At this point, the Aclar was removed from the monolayer cells samples. The samples (both monolayer and spheroids) were left to further polymerize at 60 °C for other 24 hours. The samples were sectioned with a Leica UC7 Ultramicrotome at 60 nm, silver colored on water bath. The sections were collected on copper grids (Gilder) covered with a thin formvar-film. The sections were post-stained with UA (4% UA in 50% ethanol) and 1% lead citrate in 0.1 M NaOH before microscopy. Monolayer and spheroid sections were imaged using a FEI Tecnai 12 with a tungsten filament, with an accelerating voltage of 80 kV.

The anhydrous preparation method was adapted from previous studies (19, 22). Monolayer cells were cultured for 3 and 4 weeks in both RM and OM. Samples were first washed with PBS and then incubated with ethylene glycol for 24 hours. After washing in absolute ethanol followed by acetonitrile, samples were infiltrated with increasing concentrations of epoxy resin in acetonitrile (1:1 and 3:1, 24 hours each), followed by 100% epoxy resin for 24 hours and 100% epoxy resin under vacuum (200 mbar) for an additional 24 hours. Samples were then embedded in fresh epoxy resin and polymerized at 60 °C for 24-48 hours. Sectioning, post-staining and imaging were performed following the same procedures as described for the traditional chemical fixation method.

### Scanning Transmission Electron Microscopy and Energy Dispersive X-ray Spectroscopy analysis

Highangle annular dark-field scanning transmission electron microscopy (HAADF-STEM) and energy dispersive X-ray spectroscopy (EDX) were performed using a JEOL JEM-ARM300F2 transmission electron microscope (JEOL Ltd.). The instrument was operated at an accelerating voltage of 200 kV and equipped with a cold field emission gun (CFEG) and a probe Cs corrector for STEM imaging. Elemental analysis was conducted using an energy dispersive X-ray spectroscopy detector (158 mm^2^ active area, JEOL), allowing for the detection and spatial mapping of key elements.

## RESULTS and DISCUSSION

In our previous work (13), the combination of Coherent Raman Scattering, in particular Stimulated Raman scattering (SRS), and second harmonic generation (SHG) microscopy allowed to perform a label-free identification of collagen and phosphate-rich mineral deposits in spheroids cultured for 4 weeks in osteogenic media (OM) and in monolayer cells cultured for 4 weeks in OM (Figure S1). While these approaches provide chemical specificity and spatial mapping of matrix components, they do not allow for direct visualization of ultra-structural details associated with matrix formation. This study starts from these findings, by employing transmission electron microscopy (TEM) to investigate the nano-scale organization of extracellular matrix (ECM) and to assess how sample preparation influences its preservation.

The aim of this study was to evaluate and compare two TEM sample preparation protocols, in particular conventional chemical fixation and anhydrous method, to evaluate their ability to preserve the mineralized ECM produced by MC3T3-E1 cells cultured as monolayers. In addition, the potential influence of the contrasting solutions on the morphology of the deposited mineral was assessed by imaging both contrasted and non-contrasted thin sections. Following this methodological analysis, the protocol providing the most suitable balance between matrix and structural preservation was applied to three-dimensional (3D) spheroid models. For this complex cell model, the aim was not only to assess mineral deposition, but to also examine the ultra-structural organization of the spheroids at a level of details that it is not accessible with conventional imaging approaches, such as confocal microscopy. This allowed for a detailed evaluation of the spatial relationship between cells, collagenous matrix, and mineral depositions, to determine whether mineralization within the spheroids could be considered relevant for more advanced bone studies.

Several studies have highlighted the potential limitations of conventional fixation protocols involving aqueous solutions for the study of mineralized tissues. Because these preparation methods involve multiple aqueous steps, partial dissolution or alteration of the mineral phase may occur. As a result, the morphology and distribution of mineral deposits observed by TEM may not fully reflect their native state (20, 24, 25). To address these limitations, alternative preparation strategies based on anhydrous processing have been proposed, aiming to minimize interactions between the mineral phase and aqueous reagents and improve the preservation of the inorganic component of the ECM. This anhydrous approach has been applied mostly to native bone tissue sections (20, 21, 25), whereas its use in *in vitro* culture systems has been more limited (19, 22).

### Chemical fixation for ultra-structural and mineral analysis

MC3T3-E1 subclone 4 cells were cultured on Aclar strips under regular media (RM) and OM conditions for 2 and 3 weeks. Cells cultured in RM for both 2 and 3 weeks did not exhibit detectable ECM deposition (Figure 1A, B, F and G). At both time points, only a thin cell layer was observed. The cell ultrastructure was well preserved, with intact membranes and clearly distinguishable intracellular components, such as nuclei and mitochondria, indicating that chemical fixation protocol maintained overall cellular integrity. No morphological differences were observed between samples cultured for 2 and 3 weeks under RM conditions.

**Figure 1.**
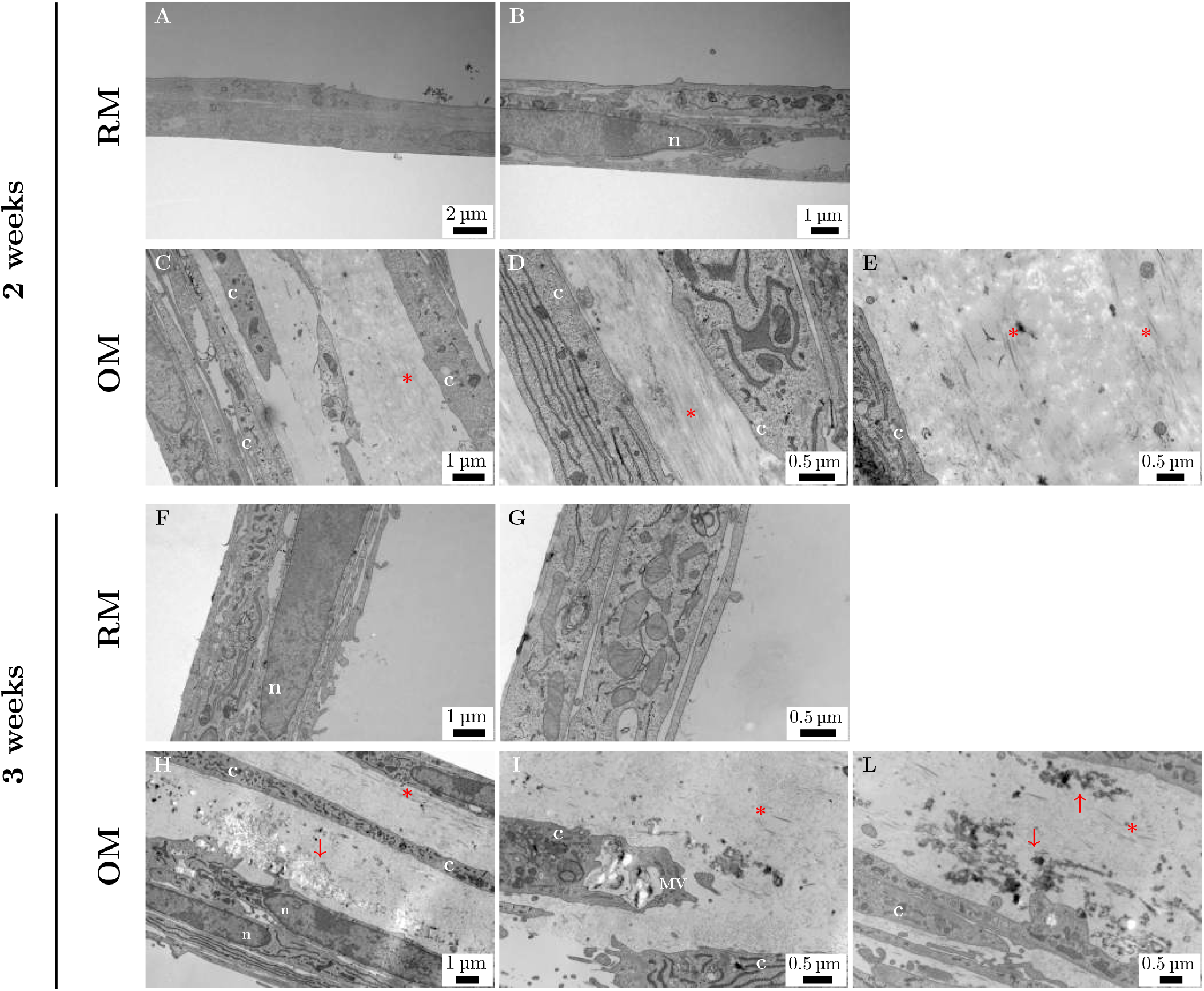
TEM images of MC3T3-E1 cells cultured as monolayers under RM and OM conditions, prepared using the chemical fixation method. A-B) Cells cultured for 2 weeks in RM. A thin cell layer is visible, with preserved intracellular organelles, including nuclei (n). C-D-E) Cells cultured for 2 weeks in OM. A thicker cell layer is observed, with cells (c) embedded within a collagenous ECM (*). F-G) Cells cultured for 3 weeks in RM. The cell layer remains thin, with clearly distinguishable organelles such as mitochondria and nuclei (n). H) Cells cultured for 3 weeks in OM,showing a thick cellular (c) and collagen-rich ECM (*) layer. Electron-dense deposits (arrow) are visible within the ECM. I) Higher magnification image of a matrix vesicle (MV) located at the peripheral region of a cell, oriented toward the ECM. Mature collagen fibrils (*) are visible in the surrounding matrix. L) Higher magnification of a region showing cells (c) and electron-dense mineral deposits (arrow) within the collagenous ECM (*).

In contrast, cells cultured in OM for 2 weeks exhibited a substantial increase in cell layer thickness and evident ECM deposition (Figure 1C). The ECM was predominantly composed of collagen fibrils, which were easily identified by their characteristic fibrillar morphology. Both highly ordered and more disorganized fibrillar arrangements were observed. In regions of aligned fibrils, the characteristic banding pattern of collagen was clearly visible (Figure 1D). ECM deposition was localized in the intercellular spaces. Despite the abundant collagen deposition, only a small amount of electrondense material was observed. The presence of these electrondense deposits in bone cell models is typically associated with matrix mineralization. At this time point, only sparse deposits were detected within the collagenous ECM (Figure 1E), suggesting that mineral formation is still at an early stage. This observation is consistent with our previous findings using second harmonic generation microscopy, where collagen-rich ECM was observed at a comparable timeline (26).

The collagenous matrix at 3 weeks maintained structural characteristics comparable to those observed at 2 weeks, with both ordered and disorganized fibrillar regions. However, at week 3, a marked increase in electron-dense structures was observed within the ECM compared to week 2 (Figure 1H and L). These structures were distributed throughout the collagenous matrix and showed heterogeneous contrast, with electron-transparent regions mixed with more electron-dense areas. These electron-dense deposits exhibited an elongated, needle-like morphology. This morphology is consistent with the elongated apatite structures reported in previous studies on matrix mineralization (16, 22, 27–30). Together with their localization within the collagenous ECM, these morphological characteristics support the interpretation that these electron-dense structures correspond to partially crystallized mineral deposits forming during osteogenic differentiation. In addition, similar electron-dense structures were also identified within intracellular vesicles located near the plasma membrane, particularly in regions oriented toward the collagen-rich ECM (Figure 1I). Their localization and morphology are consistent with matrix vesicles (MVs) described in bone cell systems and suggest that these vesicles may contain mineral precursors involved in the nucleation process (19).

To exclude the possibility that the observed electron-dense deposits resulted from staining artifacts introduced during contrast staining, additional imaging was performed on non-contrasted sections. This approach was adopted considering that contrasting solutions, in particular uranyl acetate, have a strong affinity for (calcium) phosphate and may influence the appearance of mineralized structures (31–33). Overall, no major differences were observed between contrasted and non-contrasted sections, except the slightly reduced contrast in the non-contrasted sections. In non-contrasted samples, collagen fibrillar organization was well maintained and the characteristic banding pattern was easy to observe (Figure 2 A). Importantly, comparable electron-dense structures were detected, although with slightly reduced overall contrast (Figure 2B and C). Notably, the needle-like morphology of the more electron-dense deposits was preserved, further supporting their interpretation as mineral deposits rather than artifacts introduced during the contrasting process.

**Figure 2.**
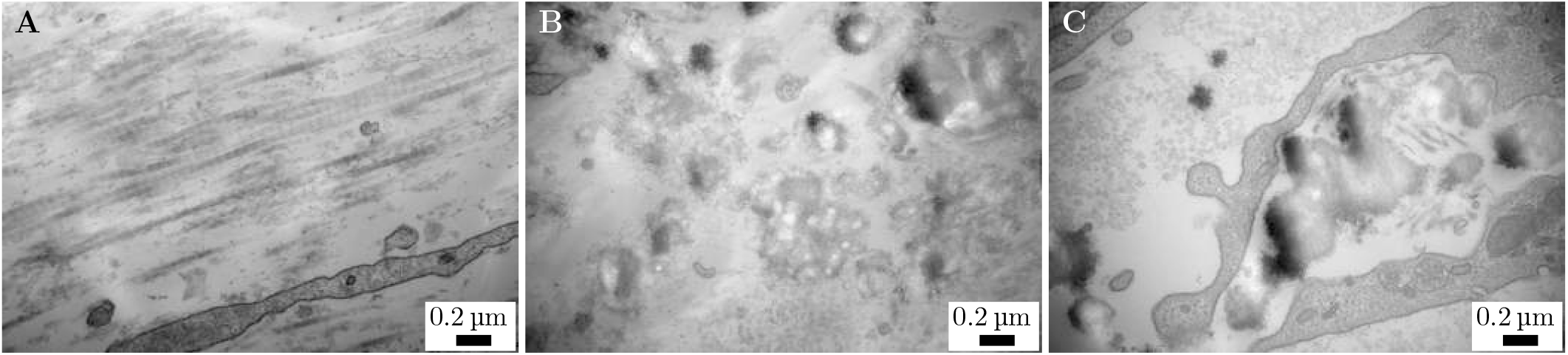
TEM images of MC3T3-E1 cells cultured as monolayers for 3 weeks in OM, obtained from non-contrasted sections prepared using the chemical fixation method. A) Mature collagen fibrils, showing the typical banding pattern. B) Deposits located in the ECM, showing differences in electron density: both more and less electron-dense regions show the typical needle-like morphology associated with mineral deposits. C) Higher magnification image of a MV, located at the peripheral region of a cell. A small aperture is visible at the lower edge of the cell membrane, suggesting that the vesicle content is being released into the extracellular environment. These deposits present the characteristic needle-like morphology typical of mineralized deposits.

To further validate these observations, and confirm that the electron-dense deposits correspond to mineral deposits, High-Angle Annular Dark-Field (HAADF) Scanning Transmission Electron Microscopy (STEM) and energy-dispersive X-ray spectroscopy (EDX) imaging and analysis were performed on the same 3-week non-contrasted samples. This approach was selected to assess the presence of phosphorous (P) and/or calcium (Ca) in the electron-dense deposits. The spectra were largely dominated by signals from elements such as uranium (U) and osmium (Os), originating from the sample preparation step, and from carbon (C) and oxygen (O), which are present both in the biological matrix and the embedding resin. A detectable P signal was observed in regions corresponding to electron-dense deposits in the collagenous matrix, whereas Ca was not detected (Figure 3). The limited detection of Ca may be attributed to several factors. The relatively low mineral content at this stage may result in signals that fall near the detection limit of this technique. However, the use of uranyl acetate during sample preparation, which includes an incubation step before embedding, may be the step influencing the preservation and detectability of the mineral phase. Given the acidic nature of uranyl acetate, partial dissolution or alteration of immature calcium phosphate deposits may occur (34, 35), especially during long incubation steps, potentially reducing the detectable Ca (and P) signal. Furthermore, overlap of signals coming from heavy metals can hide weaker elemental contributions. Despite these limitations, the detection of P in spatial association with electron-dense structures, together with their characteristic morphology and localization in the ECM, supports their interpretation as mineral deposits.

**Figure 3.**
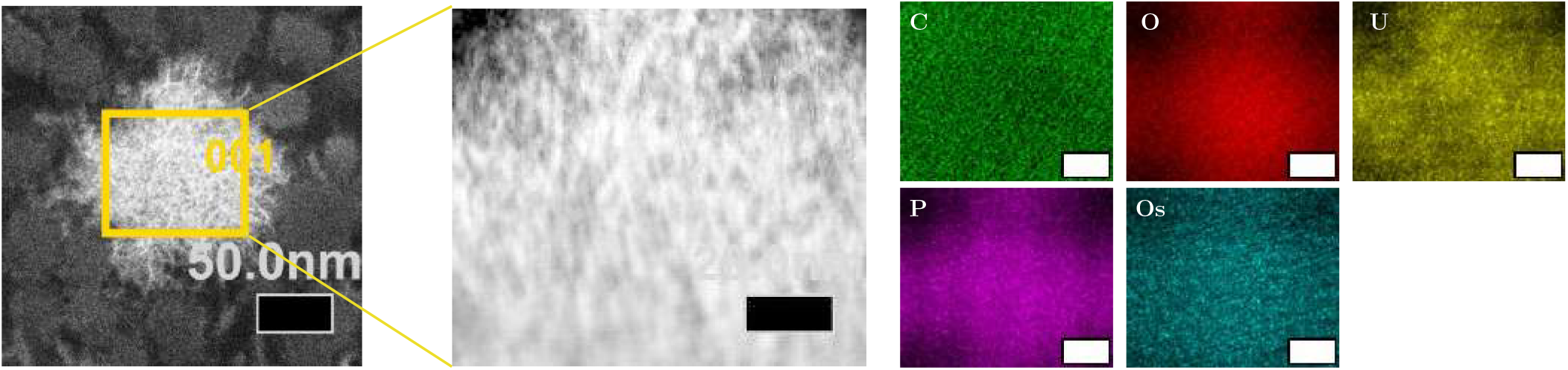
STEM-HAADF images and corresponding EDX elemental analysis of non-contrasted MC3T3-E1 monolayer cells cultured for 3 weeks in OM. The left image shows a low magnification HAADF overview of the analyzed region, which correspond to a small electron-dense deposit located in the collagenous ECM. The yellow square highlights the area selected for further EDX elemental analysis. The central image shows a higher-magnification HAADF image of the selected region. The images on the right show the corresponding EDX elemental maps, illustrating the spatial distribution of carbon, oxygen, uranium, phosphorous and osmium. Scale-bar of central and right images: 20 nm.

Additional comparison between contrasted and non-contrasted samples showed that, in both cases, some deposits exhibit heterogeneous contrast, containing regions that are more electron-transparent as well as areas that are more electron-dense. The presence of these features in non-contrasted samples suggests that these variations cannot be attributed only to the final contrasting step. As discussed above, uranyl acetate is also introduced during the earlier stages of sample preparation and may influence the preservation of immature mineral phases. So, the heterogeneous contrast observed within some deposits may reflect, in part, alterations introduced during sample preparation. However, these variations could also arise from intrinsic differences in mineral composition, density or maturation. Therefore, the observed heterogeneous contrast likely arises from a combination of intrinsic features of the mineral phase and possible alterations introduced during sample preparation, in particular by uranyl acetate incubation.

Moreover, comparable electron-dense structures were not observed in the 2 weeks OM samples, further supporting the interpretation that the structures detected at week 3 correspond to newly formed mineral deposits associated with ECM mineralization, and are not originated from uranyl acetate or other heavy metals used during sample preparation (as RM and OM samples were prepared in the same way).

### Anhydrous method for analysis on mineral preservation in monolayer cell cultures

Based on the limited mineral deposition observed at week 2 using the chemical fixation method, the anhydrous preparation protocol was applied to samples cultured for 3 and 4 weeks. Cells were maintained under the same culture conditions as previously described and grown on the same Aclar substrate to ensure comparability between the samples.

As expected, this preparation approach did not optimally preserve cellular ultrastructure as it is primarily intended to retain the ECM and mineral phase rather than intracellular components.

Samples cultured in RM from 3 and 4 weeks exhibited only a very thin layer of material, which can be attributed to cells (Figure 4A,B, F and G). The observed structure lacked the well-preserved morphology seen with chemical fixation and appeared largely disrupted. The residual material resembled cellular debris rather than intact cells, consistent with the limited preservation capacity of this method for cellular components. No evidence of ECM deposition or minerallike electron-dense structures was detected under these conditions.

**Figure 4.**
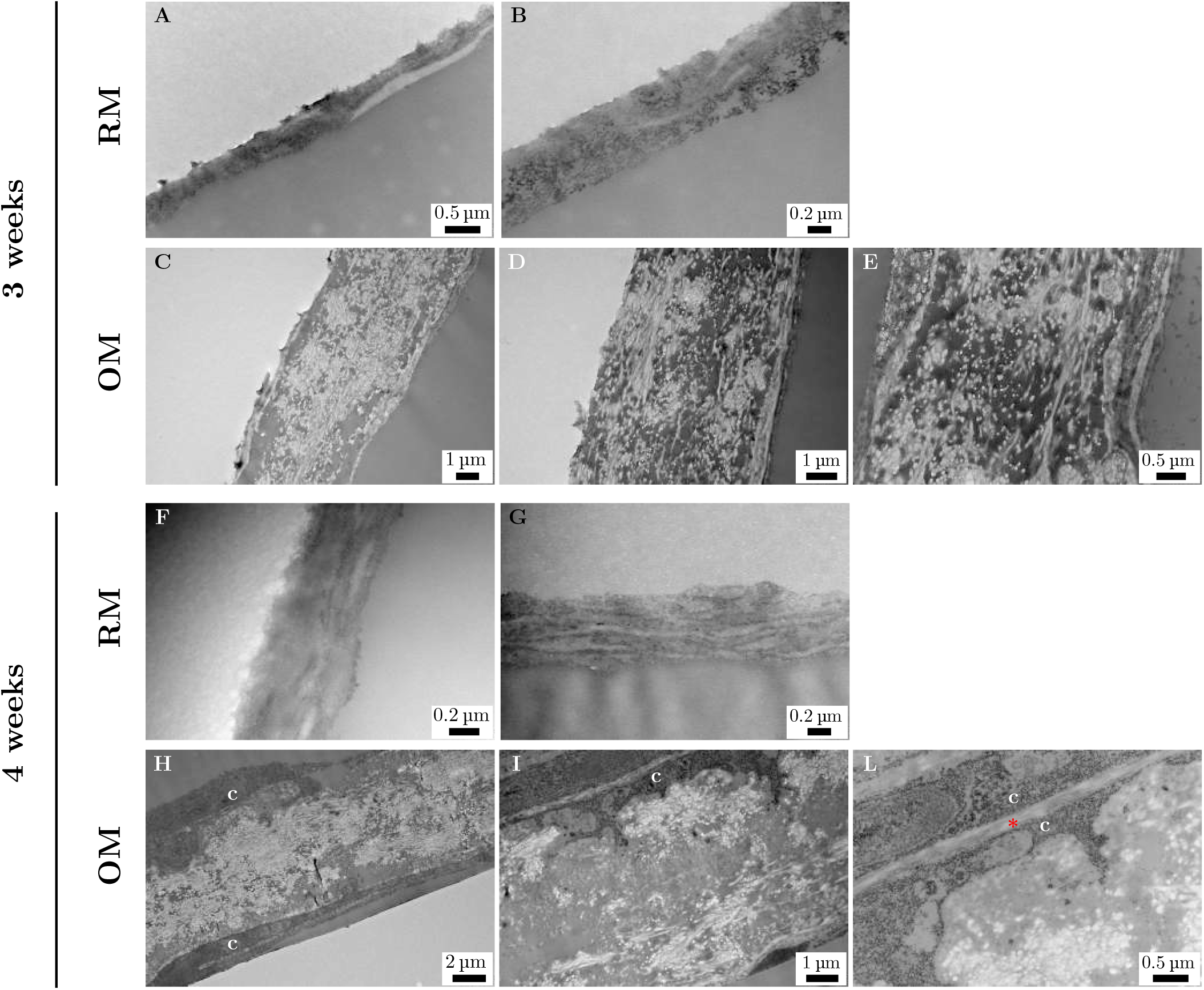
TEM images of MC3T3-E1 cells cultured as monolayer cells under RM and OM conditions and prepared using the anhydrous method. A-B) Cells cultured for 3 weeks in RM. A thin residual layer of material is observed, likely corresponding to cellular debris. No distinct cellular or extracellular structures can be identified. C-D-E) Cells cultured for 3 weeks in OM. A low-contrast ECM with a fibrillar-like organization is visible, consistent with collagen deposition. Electron-dense regions are present within the matrix. F-G) Cells cultured for 4 weeks in RM. A similar layer of cell debris is observed as for week 3, with no distinguishable cellular or ECM structures. H-I) Cells cultured for 4 weeks in OM. Partially preserved cell profiles (c) can be identified. The surrounding ECM appears more extensive and contains numerous electron-dense regions. L) High magnification image showing two cell profiles (c) and collagen fibrills (*) displaying a recognizable fibrillar organization. Electron-dense regions are also observed in the ECM.

In contrast, samples cultured in OM for 3 weeks displayed a distinguishable ECM region characterized by a low-contrast, fibrillar-like network (Figure 4C, D and E). Although individual fibrils were not clearly distinguishable, the matrix displayed a faint, striated organization, suggesting that this area could correspond to the deposited collagen fibrils. The reduced contrast of these fibrillar structures indicates that the anhydrous preparation method may compromise the preservation or visualization of the organic matrix. Within this matrix, electron-dense structures were observed. Morphologically, these electron-dense deposits appeared more compact and homogeneous, lacking the internal contrast variation and needle-like features observed in chemically fixed samples.

At 4 weeks in OM, a marked increase in the abundance and spatial distribution of both fibrillar ECM and electron-dense structures were observed compared to the 3-week samples (Figure 4H, I and L). Based on these first observations alone, the increase in the electron-dense material could be interpreted as progressive mineral deposition, consistent with previous observations indicating that the majority of mineral deposition occurs between week 3 and 4 (13).

Despite the known limitations of the anhydrous method for cellular preservation, some cell profiles were partially recognizable in week 4 samples (Figure 4I and L). Although ultra-structural details were not well maintained, general cell morphology and in some instances nuclear regions could be distinguished. Additionally, some collagen fibrils appeared well preserved, with discernible fibrillar organization (Figure 4L). This improved matrix visibility may be related to increased matrix thickness and compaction at later stages of culture, which may facilitate retention of structural features even un-der conditions that are less favorable for cellular preservation. To further investigate the nature of the observed fibrillar and electron-dense regions, non-contrasted samples cultured for 4 weeks in OM were analyzed, in combination with STEM-HAADF imaging and EDX spectroscopy.

Imaging of non-contrasted samples revealed no major differences in ECM morphology or organization compared to contrasted samples (Figure 5 A, B and C), indicating that the contrasting step does not significantly alter the structural appearance of the ECM components in anhydrous preparations. This observation supports the interpretation that the structural alterations and reduced mineral preservation observed in chemically fixed samples are not caused by the contrasting step itself, but by the cumulative effects of aqueous processing and the uranyl acetate incubation step prior to embedding. EDX analysis provided further insight into the elemental distribution within the samples. In regions corresponding to fibrillar ECM, spectra were dominated by C and O, as expected from the organic matrix and embedding resin (Figure 6). A detectable P signal was observed in these fibrillar regions, whereas Ca was not detected (Figure 6). In contrast, electron-dense regions did not exhibit a measurable P signal and showed spectra comparable to those obtained from the surrounding epoxy resin. Trace of silicon (Si) signal were also detected and are likely attributable to contamination from glassware used during sample preparation.

**Figure 5.**
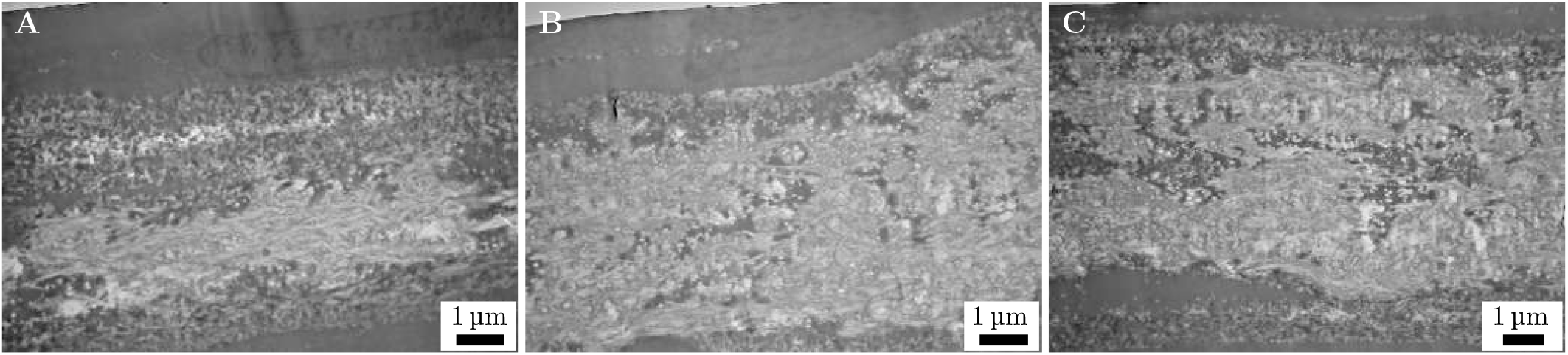
TEM images of non-contrasted sections of MC3T3-E1 monolayer cells cultured for 4 weeks in OM and prepared using the anhydrous method. In all images, a low-contrast collagenous ECM with a fibrillar morphology is observed, consistent with collagen organization. Electron-dense regions are present within the matrix, with no apparent difference in morphology compared to contrasted samples. A-B) The outline of a cell nucleus can be distinguished within the surrounding ECM.

**Figure 6.**
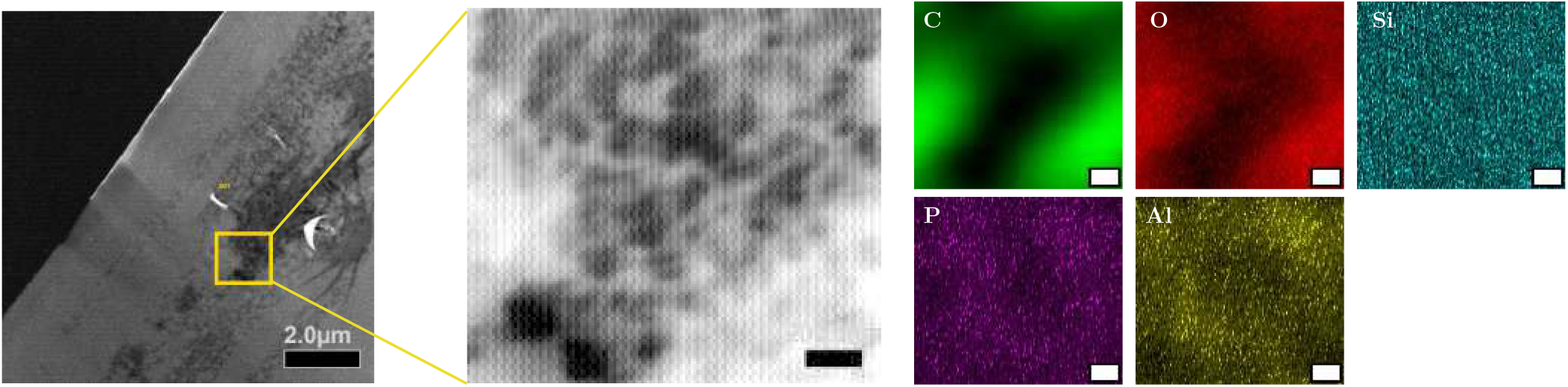
STEM-HAADF images and corresponding EDX elemental analysis of non-contrasted MC3T3-E1 monolayer cells cultured for 4 weeks in OM and prepared using the anhydrous method. The left image shows a low-magnification HAADF overview of the analyzed region, corresponding to a fibrillar area consistent with ECM organization. The yellow square highlights the area selected for further EDX elemental analysis. The central image shows a higher-magnification HAADF image of the selected region. The right panels show the corresponding EDX elemental maps, illustrating the spatial distribution of carbon (C), oxygen (O), silicon (Si), phosphorous (P) and aluminum (Al). Scale-bar of central and right images: 20 nm.

These findings suggest that the electron-dense regions observed in anhydrous samples may not directly correspond to phosphate-rich mineral deposits. Instead, they may originate from resin accumulation or preparation-related artifacts, potentially occupying spaces previously associated with mineral phases prior to processing. The presence of the P within the fibrillar ECM however indicates that P-containing species are associated with the organic matrix, which is consistent with early stages of mineralization occurring within or along collagen fibrils.

Taken together these results indicate that while the anhydrous method allows for visualization of ECM organization, it may not reliably preserve or represent the mineral phase in cell culture samples. Although electron-dense structures increase over time and are observed exclusively under osteogenic conditions, their lack of elemental confirmation suggests that caution is required when interpreting them as mineral deposits.

Overall, while this method has been successfully used for native mineralized tissue, its suitability for *in vitro* cell culture systems appears more limited. The combined ultra-structural and compositional analysis suggests that the anhydrous protocol may introduce artifacts or alter the spatial distribution of mineral phases, and therefore should be applied with care when investigating mineralization in such models.

### Spheroids ultra-structural analysis via chemical fixation

For spheroids characterization, preservation of both spheroids ultrastructure and ECM organization is essential to evaluate the spatial relationship between cells and matrix components. For this reason, the conventional chemical fixation protocol was applied to spheroids cultured for 5 and 6 weeks. It was decided to analyze spheroids at this time point because from our previous studies we observed that ECM mineralization happens at a slightly slower rate in spheroids compared to monolayer cell cultures (13, 26).

Based on our previous findings demonstrating that spheroids cultured in RM do not produce collagenous ECM and become structurally unstable after 3 weeks (26), RM samples were not systematically analyzed at later time points. At 6 weeks, RM spheroids (observed with bright field microscopy and confocal laser scanning microscopy) appeared structurally compromised with collapse of the aggregate, preventing reliable ultra-structural evaluation.

Several spheroids cultured in OM for 5 weeks were imaged. Overall, the spherical architecture was well preserved, maintaining a defined and compact 3D structure (Figure 7A). A collagen-rich ECM was clearly visible, with mature fibrillar organization. Both highly organized and less ordered collagen-rich regions were observed. In one representative spheroid, collagen deposition was predominantly localized in the central region of the aggregate. The fibrillar structure was well maintained, with clearly distinguishable collagen fibers (Figure 7B and C). Electron-dense deposits were on the other hand not visible in this sample. In another representative spheroid, electron-dense deposits within the ECM were more evident (Figure 7D and F). In this case, deposition appears more prominent in the outer layer of the aggregate. This spheroid was notably larger than others in size, with abundant cells and a well-organized collagen network (Figure 7D and E). The electron-dense deposits displayed a needle-like morphology comparable to those observed in monolayer cultures prepared using the same chemical fixation method. Non-contrasted images did not show any differences in the morphology of the collagenous matrix compared to contrasted samples, only a slightly reduced contrast (Figure 8A and B. Figure 8A was taken in the same position as Figure 7C). Electron-dense deposits were also observed in the non-contrasted sections, with needle-like morphology comparable to contrasted spheroids and monolayers, supporting their identification as mineral deposits (Figure 8C). No direct interaction between the spheroids and the surrounding alginate hydrogel was observed, and presence of debris was observed in the alginate pocket.

**Figure 7.**
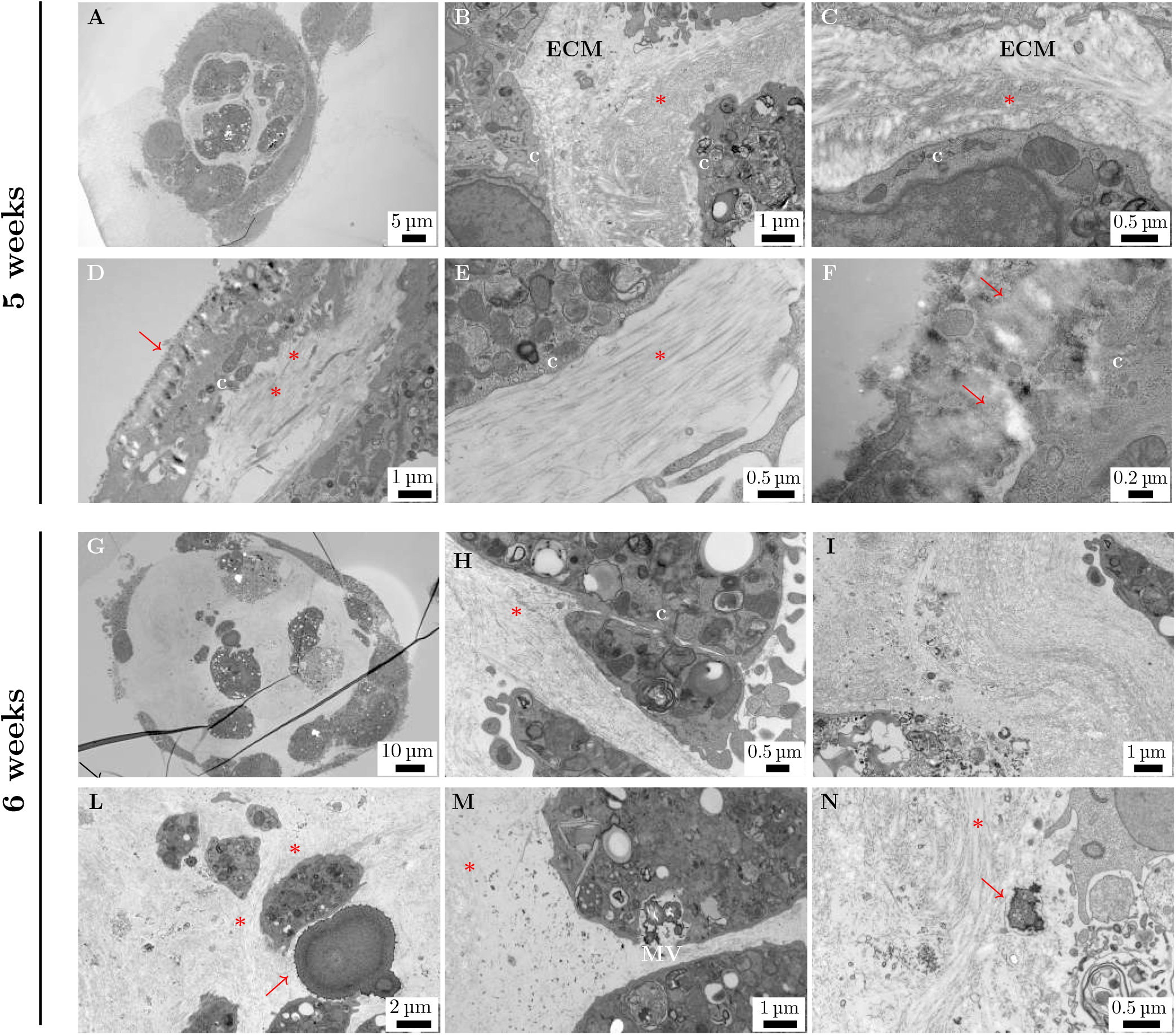
TEM images of alginate-encapsulated MC3T3-E1 spheroids cultured in OM and prepared using the chemical fixation method. A-F) Spheroids cultured for 5 weeks in OM. A) Low magnification image showing the overall spheroids organization. B-C) High magnification images of distinct regions showing small, less organized collagen fibrils (*), extracellular matrix (ECM) and cells (c). D) Peripheral region of a spheroids showing both organized, mature collagen fibrils and electron-dense structures. E) High magnification image showing an area with mature and highly organized collagen fibrils. F) High magnification image of electron-dense deposits, showing the characteristic needle-like morphology typical of mineral deposits. G-N) Spheroids cultured for 6 weeks in OM. G) Low magnification image showing the overall spheroids organization. H) High magnification image of an area of organized collagen fibrils (*) and cells (c). I) High magnification image of an area of extensive deposited and organized collagen fibrils. L) High magnification image of an area of the ECM showing the collagenous matrix and an electron-dense deposit. M) High magnification image of a MV, containing electron-dense structures, opening towards the extracellular environment. N) High magnification image of an area of the ECM showing an organized collagenous matrix and an electron-dense deposit, with the typical needle-like morphology of mineral deposits.

**Figure 8.**
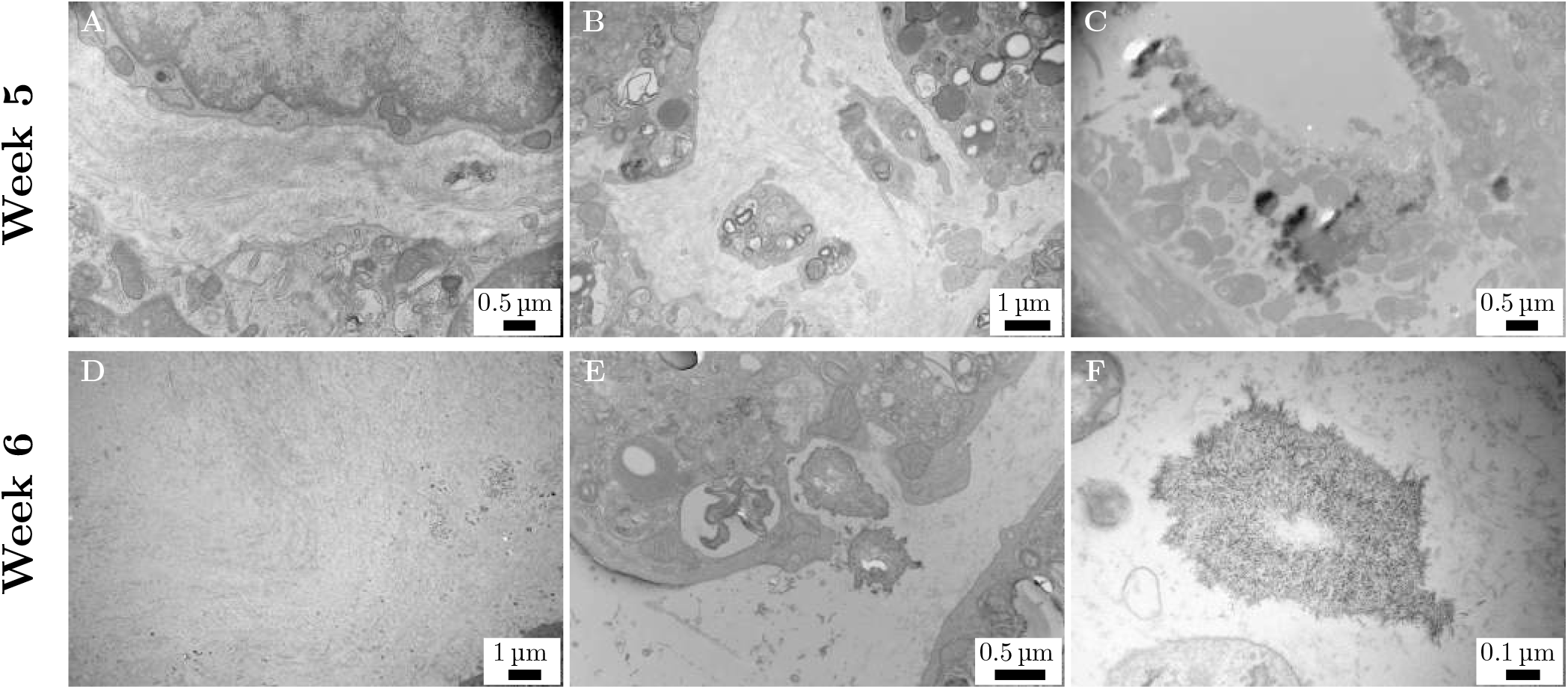
TEM images of alginate-encapsulated MC3T3-E1 spheroids cultured in OM, obtained from non-contrasted sections and prepared using the chemical fixation method. A) High magnification image of a region of the spheroid, showing small, less organized collagen fibrils. B) High magnification image of a region of the spheroid, showing mature, more organized collagen fibrils. C) High magnification image showing electron-dense deposits with the characteristic needle-like morphology of mineralized structures. D) High magnification image of an area of extensive deposited and organized collagen fibrils. E) High magnification image of a MV opening toward the extracellular environment and releasing its contents; the electron-dense material present a needle-like morphology. F) High magnification image of an electron-dense deposit, showing the typical needle-like morphology of mineral deposits.

Spheroids cultured for 6 weeks in OM exhibited increased collagen deposition compared to week 5 (Figure 7G). The aggregate maintained a well-defined spherical shape, with a dense and organized fibrillar ECM. Collagen fibers appear aligned and structurally mature (Figure 7H and I). Within the collagenous matrix, small electron-dense deposits were observed (Figure 7L and N). Their morphology and localization were consistent with the mineral-like structures previously described in monolayer cell cultures. In several regions, these deposits displayed an elongated, needle-like morphology within the fibrillar ECM, further supporting their identification as mineral deposits. Additionally, MVs were detected in proximity to the collagen matrix, and appeared to open toward the extracellular space (Figure 7M), a characteristic consistent with the vesicle-mediated nucleation process described in mineralizing tissues (28, 36, 37). No differences in collagen organization and structure were observed in the non-contrasted sample (Figure 8D). Importantly, similar electron-dense structures were observed in non-contrasted sections (Figure 8E and F), further confirming that these features do not arise from staining artifacts but represent intrinsic components of the mineral matrix.

To further support this observation, STEM-HAADF and EDX imaging and analysis was performed on non-contrasted 6-week cultured spheroids sections to assess the presence of P and/or Ca within these electron-dense deposits. Although Ca was not reliably detected (only in one analyzed area, data not shown), likely due to the low mineral content and signal overlap with the heavy metals, P was repeatedly detected in spatial association with the electron-dense deposits located in the ECM (Figure 9A) and the observed MV (Figure 9B), providing additional support for the identification of these regions as mineral deposits.

**Figure 9.**
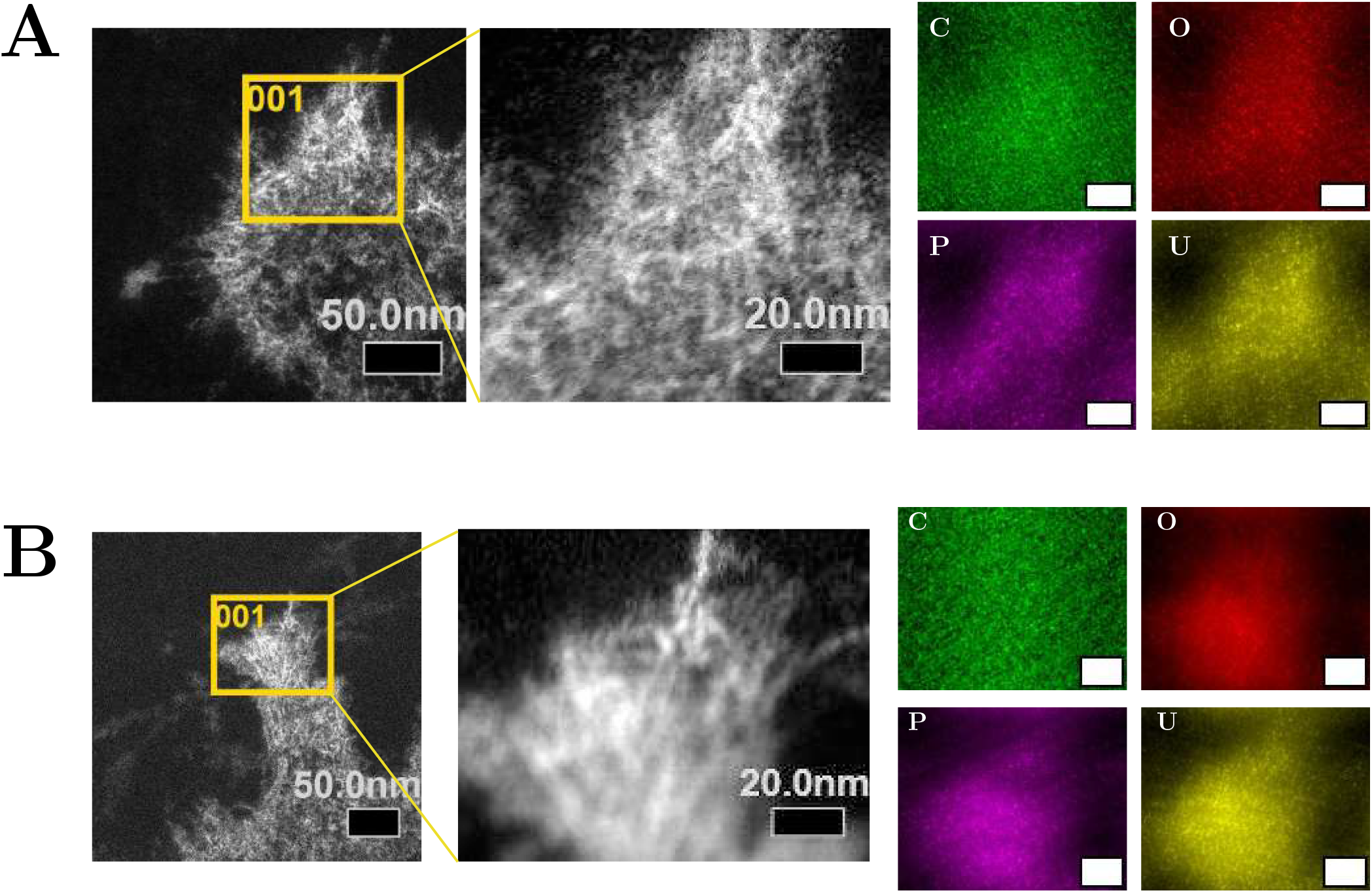
STEM-HAADF images and EDX elemental analysis of electron-dense structures in non-contrasted spheroids cultured for 6 weeks in OM. (A) Electron-dense deposit located within the collagenous ECM. The left image shows a low magnification HAADF overview of the analyzed region. The yellow square highlights the area selected for further EDX elemental analysis. The central image shows a higher-magnification HAADF image of the selected region. The images on the right show the corresponding EDX elemental maps, illustrating the spatial distribution of carbon, oxygen, phosphorous and uranium. (B) Electron-dense deposits located in a MV opening up and releasing its contents into the collagenous ECM (see Figure 8E). The left image shows a low magnification HAADF overview of the analyzed region. The yellow square highlights the area selected for further EDX elemental analysis. The central image shows a higher-magnification HAADF image of the selected region. The images on the right show the corresponding EDX elemental maps, illustrating the spatial distribution of carbon, oxygen, phosphorous and uranium. Scale-bar of right images in A and B: 20 nm.

Interestingly, differences in the internal contrast of the electron-dense deposits were observed, depending on their localization within the spheroids. Deposits located in the outer regions of the aggregates often displayed heterogeneous contrast with alternating electron-transparent and electrondense regions, similar to what was observed in monolayer cultures. In contrast, deposits detected in the central region of the spheroids, well embedded in the collagenous matrix, generally appeared more homogeneous. These observations may suggest partial alteration of the mineral phase during sample preparation. Deposits located near the surface of the spheroids are likely more exposed to staining and washing steps, while those in the core may be partially protected by the surrounding 3D matrix. This reduced exposure may limit the potential alterations of the mineral phase and contribute to the more uniform appearance of the mature deposits observed in the central region of the spheroids.

Although mineral-like electron-dense deposits were detected within the spheroids ECM, their overall abundance appeared lower compared to monolayer cultures at comparable differentiation time-points, as well as to observations reported in our previous study, where Stimulated Raman Scattering microscopy was used to chemically identify phosphate deposits in 4-weeks old spheroids (13). This difference may reflect both biological and technical factors. From a biological perspective, mineralization within 3D aggregates may proceed more gradually due to differences in matrix organization, cell density, and diffusion of nutrients or ions within the spheroid structures. In addition, mechano-biological cues may contribute to these differences. Additionally, variability in mineralization timing between spheroid batches may arise from differences in spheroid size, cell passage number, number of cells making up the spheroids and other culture-related parameters. From a technical point of view, early mineral nuclei forming within and in between the collagen matrix are expected to be small and poorly crystalline during the initial stages of mineralization. These immature deposits may be particularly sensitive to the aqueous steps and uranyl acetate incubation used in the chemical fixation method. As a consequence, small mineral precursors located within or between collagen fibrills may be partially dissolved or displaced during sample preparation, potentially leading to an underestimation of the total amount of mineral present in the spheroid ECM.

Overall, the presence of collagen-rich ECM, MVs, and needle-like electron-dense deposits, together with the detection of P within these regions, supports the interpretation that mineralization occurs within this 3D culture model and follow ultra-structural features consistent with those observed in osteogenic monolayer cultures. These findings are in line with results obtained in 2D cultures of primary cells (17), where electron-dense mineral deposits associated with matrix vesicles and surrounding lacuna-like structures were described. In their study, they also noted the presence of uranyl acetate staining, attributed to its interaction with negatively charged, mineral-binding molecules. In comparison with these observations, our results suggests that similar mechanism of MVs-mediated mineral nucleation and associated ultra-structural features are maintained in 3D spheroid model.

## CONCLUSIONS

In this study, we compared conventional chemical fixation and anhydrous preparation methods for transmission electron microscopy (TEM) to evaluate their ability to preserve the mineralized extracellular matrix (ECM) produced by MC3T3-E1 subclone 4 cells in monolayer cell cultures and as alginate-encapsulated spheroids.

Chemical fixation preserved cellular morphology and collagen organization, allowing a detailed analysis of the cell-ECM interactions. Imaging of contrasted and non-contrasted sections revealed that uranyl acetate and aqueous solutions used during sample preparation may alter immature mineral phases, compromising accurate visualization of early mineral deposits. Uranyl acetate is introduced at two points during sample preparation, first after dehydration and before resin embedding, and later as contrasting agent before imaging. Although comparison between contrasted and non-contrasted sections revealed no major differences aside from image contrast, comparison with previously published results (13) revealed a reduced amount of mineral deposits in the present samples. This suggests that the incubation step prior to embedding might be the source of mineral alteration. This interpenetration is also further supported by HAADF-STEM data, where uranyl is detected among the present elements in mineralized regions.

In contrast, the anhydrous preparation method provided limited preservation of cellular ultra-structure and altered the appearance of ECM components. Elemental analysis indicated that phosphorous was associated with fibrillar regions, while electron-dense areas lacked a corresponding signal, suggesting that these features may not reliably represent mineral deposits. Since this protocol does not involve aqueous solutions and uranyl acetate is used only as contrasting agent, the reduced detection of mineral deposits reflects intrinsic limitations of the anhydrous method, making it overall less reliable for mineral detection compared to the traditional chemical fixation method. In addition, as for the chemical fixed samples, no major differences were observed between contrasted and non-contrasted sections, further supporting the interpretation that the uranyl acetate incubation step in the chemical fixation protocol is the one influencing the state of the mineral deposits.

When applied to alginate-encapsulated spheroids, chemical fixation allowed for preservation of the overall aggregate architecture, allowing for assessment of overall structure and ECM organization and mineral deposition. These observa-tions confirm that spheroids produce a collagen-rich matrix and exhibit ultrastructure features consistent with mineralization in monolayer cell models, supporting their relevance as 3D *in vitro* models for bone research. However, the amount of mineral deposits was lower compared to previous studies and to what was detected in the same system in hydrated samples using optical microscopy (13). This difference could result from the effects of uranyl acetate and aqueous solutions (as observed for monolayer cells) used during sample preparation, but also from potential biological factors, such as spheroids size, cell passage, and maturity state of the mineral deposits (small mineral precursors compared to more mature mineralized deposits).

Overall, these results demonstrate that the choice of the sample preparation method should be guided by the specific aim of the study. Chemical fixation remains the most suitable approach for preserving cellular and matrix ultra-structure, whereas the anhydrous method, in its current state, provides limited and potentially ambiguous information regarding mineral identification in cell culture models.

These findings highlight the need for improved preparation strategies that minimize chemical alteration of the mineral phase and enable more reliable correlation between ultrastructural and compositional analyses, ultimately improving the characterization of early mineralization processes.

## ACKNOWLEDGEMENTS

We acknowledge Nan Tostrup Skogaker and the Electron Microscopy lab at the Department of Clinical and Molecular Medicine, NTNU, for training, technical assistance and access to the electron microscopy core facility. We acknowledge Stiftelsen Biopolymer for financial support. We acknowledge Dr. Alexandra Porter for providing the protocol for the anhydrous sample preparation method. We acknowledge Sebastian J Kjeldgaard-Nintemann and Astrid Bjørkøy for their help and support with CRS experiments. The authors would like to acknowledge support from the Research Council of Norway through the Norwegian Center for Transmission Electron Microscopy, NORTEM (197405).

