## Supplementary info for "Ultra-structural analysis of mineralized extracellular matrix in osteogenic monolayers and spheroids: comparison of sample preparation methods"

### Supplementary Section 1 Material and methods

#### Coherent Raman Scattering

Coherent Raman Scattering (CRS) microscopy and second harmonic generation (SHG) microscopy were performed on an inverted Leica Stellaris 8 CRS microscope (Leica Microsystems), equipped with a picoEmerald S dual beam infrared laser (APE). The systems provides two syncronized pulsed beams: a fixed Stokes beam at 1032 nm and a tunable pump beam from 720 to 980 nm. Stimulated Raman Scattering (SRS) was acquired in the forward (transmission) direction as energy transfer between the modulated Stokes beam and the pump beam by a photodiode. The SHG signals was detected in the reflected (epi) direction via photon-counting detectors (HyD, Leica Microsystems). The emission bandpass filter of 465/170 nm was used for SHG detection. Imaging was performed with a 25x water immersion objective (HC FLUOTAR L 25x/0.95 W VISIR, Leica Microsystems). Samples (both monolayer cell cultures and alginate-embedded spheroids) were imaged between two glass cover-slips, and the condenser was aligned for Köhler illumination. SRS imaging targeted specific vibrational modes. A pump wavelength of 938.7 nm (corresponding to a Raman shift of 960 $cm^{-1}$) was used to probe the phosphate groups, while 796.8 nm (2857 $cm^{-1}$) was used for lipid detection.

As for sample preparation, monolayer cells cultured for 4 weeks in OM were first rinsed with PBS, fixed using 4% PFA for 20 minutes and then rinsed with PBS three times for 5 minutes. Alginate-embedded spheroids cultured for 4 weeks in OM were first washed with 5 mM BaCl for 5 minutes, and then fixed using 4% PFA for 20 minutes, and rinsed three times with PBS for 5 minutes.

### Supplementary Section 2 Results and Discussion

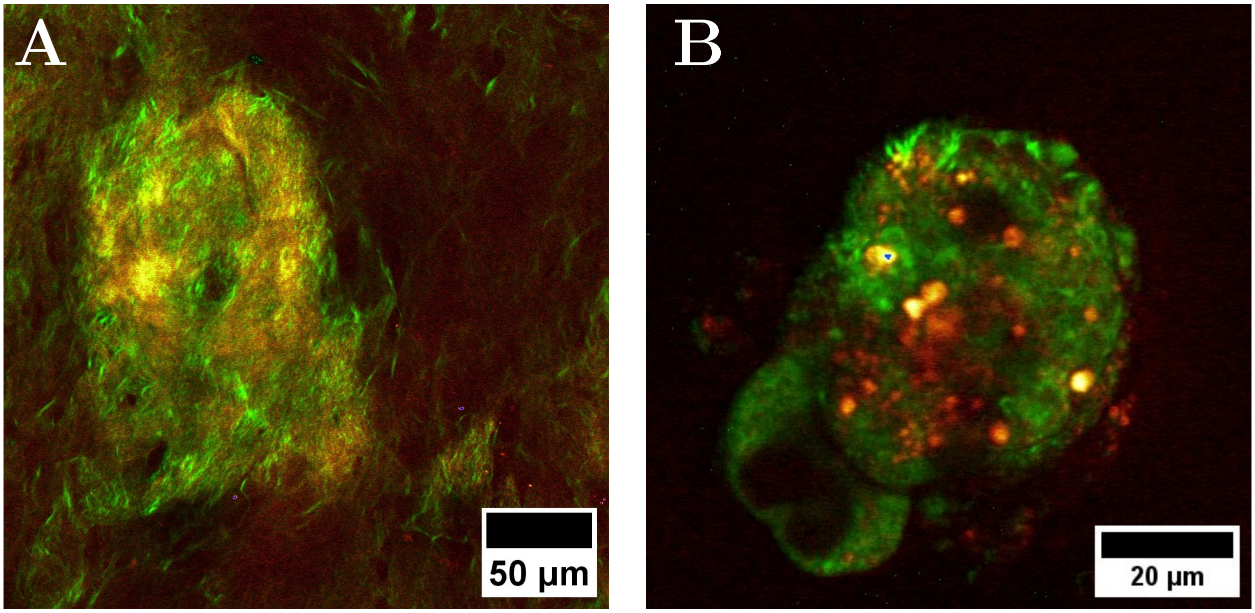


Figure S1: SRS and epi-SHG images of mineral deposits (red) and collagenous matrix (green) deposited by MC3T3-E1 subclone 4 cells cultured for 4 weeks in OM as monolayers (A) and alginate-encapsulated spheroids (B).
